# Reconstructing meaning from bits of information

**DOI:** 10.1101/401380

**Authors:** Sasa L. Kivisaari, Marijn van Vliet, Annika Hultén, Tiina Lindh-Knuutila, Ali Faisal, Riitta Salmelin

## Abstract

We can easily identify a dog merely by the sound of barking or an orange by its citrus scent. In this work, we study the neural underpinnings of how the brain combines bits of information into meaningful object representations. Modern theories of semantics posit that the meaning of words can be decomposed into a unique combination of individual semantic features (e.g., “barks”, “has citrus scent”). Here, participants received clues of individual objects in form of three isolated semantic features, given as verbal descriptions. We used machine-learning-based neural decoding to learn a mapping between individual semantic features and BOLD activation patterns. We discovered that the recorded brain patterns were best decoded using a combination of not only the three semantic features that were presented as clues, but a far richer set of semantic features typically linked to the target object. We conclude that our experimental protocol allowed us to observe how fragmented information is combined into a complete semantic representation of an object and suggest neuroanatomical underpinnings for this process.

---

The brain binds available information about objects with prior knowledge, thus allowing us to make sense of the world around us. The ability to use available information about an object (e.g. the observation of something that *has legs, is gray* and *has a trunk*), to activate relevant existing knowledge in the semantic system (e.g. *is endangered, has white tusks*) can be characterized as a process of pattern completion where few elements serve to activate a number of relevant elements in the same representation. While one can easily demonstrate the existence of such a process behaviorally, as in the example above, neuroimaging evidence of pattern completion of semantic information is critically lacking. As such, we also do not understand the neuroanatomical bases of this process. Thus, in this study, we ask whether semantic pattern completion can be demonstrated in the human brain and what brain regions are involved in integrating features together into rich representations of objects.

Many neuro-cognitive accounts on the semantic system propose that the meaning of objects can be formalized using smaller components called features (e.g. Cree et al., 2006; McRae, Sa, et al., 1997; Tyler, Moss, et al., 2000; Vigliocco et al., 2004). The features which make an object are putatively coded in a distributed fashion, primarily in the same regions that are involved in processing and perceiving them (McClelland and Rogers, 2003; Plaut and Shallice, 1993; Tyler, Moss, et al., 2000; Tyler and Moss, 2001). According to this view, a neural representation of the underlying object would be defined as a specific and relatively stable pattern of activation across the relevant feature nodes (Masson, 1995; McRae, Sa, et al., 1997; Patterson et al., 2007; Taylor, Devereux, et al., 2011; Vigliocco et al., 2004). Computational models further postulate that the activation of a sufficient number of semantic features may lead to activation of the whole semantic representation via a pattern-completion-like process (Masson, 1995; McClelland and Rogers, 2003; Plaut and Shallice, 1993). While pattern completion has been demonstrated in the visual domain (Tang et al., 2014) and in the context of episodic memory (e.g. O’Reilly and McClelland, 1994; O’Reilly and Rudy, 2001), there is little we know about the neural underpinnings of reconstructing semantic representations. In this study, we assess pattern completion of semantic information in the human brain, by making use of a multi-dimensional semantic space. Each dimension in the semantic space corresponds to a single semantic feature. The meaning of an object is defined as a position in this space (a semantic coordinate), which in turn is determined by the weighted combination of the dimensions. The distance (e.g. cosine) between two concepts quantifies their semantic similarity. Semantic spaces can be obtained, for example by using statistical co-occurrence information collected from large text corpora (Erk, 2012; Kanerva and Ginter, 2014; Mikolov, Sutskever, et al., 2013; Turney and Pantel, 2010) or using behavioral methods to estimate similarity of descriptive content between items (Devereux et al., 2014; McRae, Cree, et al., 2005; Vinson and Vigliocco, 2008; Sudre et al., 2012). Such semantic spaces have been used as priors in machine learning based neural decoding models that have successfully associated various semantic feature sets (i.e. sets of dimensions that span the semantic space) with neural signatures and, by combining them together, predicted neural activation patterns for novel objects (Broderick et al., 2018; Huth, Nishimoto, et al., 2012; Huth, Heer, et al., 2016; Just et al., 2010; Mitchell et al., 2008; Pereira et al., 2018; Simanova et al., 2010; Sudre et al., 2012). This demonstrates that the feature-based model of the semantic system is useful in describing the neural representation of meaningful stimuli.

The visual object processing system may provide insights into the neuroanatomical underpinnings of the semantic pattern completion process. Visual pattern completion has been suggested to take place in the ventral stream via recurrent connections (Tang et al., 2014; Clarke, Taylor, et al., 2011). Particularly, the perirhinal cortex (PRC), which is located in the anterior apex of this hierarchical system, has been deemed relevant in fine-grained visual analysis of objects (Barense et al., 2010; Buckley and Gaffan, 2006) and binding visual information with information from other sensory modalities (Taylor, Moss, et al., 2006; Taylor, Stamatakis, et al., 2009), including information about object meaning (Liu and Richmond, 2000). This region has been suggested to be sensitive to object-specific semantic information (Clarke and Tyler, 2014; Kivisaari, Tyler, et al., 2012). Therefore, we hypothesize that the ventral stream system, and the PRC in particular, may be involved in the pattern completion process where fragmental semantic information is completed to form a coherent object.

In this study, we probe target objects with a small set of verbal semantic features and thereby putatively facilitate activation in a rich network of other semantic features that are not presented but are related to the target object. Specifically, we mimic a guessing game where the participant is presented with a sequence of three clues (henceforth, a “clue triplet”; e.g., “has legs”, “has a thick skin”, “has a trunk”) and asked to guess the object that the clues describe (i.e., “an elephant”). We take advantage of functional magnetic resonance imaging (fMRI) and evaluate whether the bloodoxygen-level dependent (BOLD) response is best predicted by the semantic coordinates of the explicitly presented clues or a larger set of features, extending to those that were never presented to the participant (e.g., “is endangered”, “is heavy”, “does trumpet”). We hypothesize that the brain automatically ties together the presented clues with other features linked to the target object. If so, the best decoding performance would be achieved by using an even larger set of features that are associated with the target object, as compared to the exact selection of features that were presented to the participants.

The semantic space in this study is built from a large Internet-derived Finnish text corpus (Kanerva and Ginter, 2014) using the word2vec algorithm (Mikolov, Sutskever, et al., 2013; Mikolov, Chen, et al., 2013). In order to predict the brain activity to a given object/feature, we use a linear-regression decoding approach which, for each target object or semantic feature, maps its coordinates in a multi-dimensional semantic space to a corresponding BOLD activation pattern (Mitchell et al., 2008; Palatucci et al., 2009). A leave-two-out scheme is used to assess the performance of this mapping. We further employ representational similarity analysis (RSA, Kriegeskorte et al., 2008) to visualize the brain regions which are involved in representing the target objects or in completing the patterns of object features into target concepts.

## Results

### Brain activation patterns reliably predict the target objects

The neural representations of each target object were probed in the fMRI task using six different sets of clue triplets. Over all trials, the participants guessed the implied identities of the target objects at a high accuracy (93.3 percent correct; see also Table S1).

In the first analysis, we tested whether we can use the corpus-derived semantic coordinates of the target objects to decode the BOLD activation patterns elicited using clues. In order to optimize the decoding accuracy, we averaged the BOLD activation maps of the six trials for each target object. Furthermore, we restricted the analyses to a subset of voxels (*n* = 500) that showed a consistent activation pattern across the six trials (i.e., stability selection; cf. Just et al., 2010; Mitchell et al., 2008). The measurement and analysis protocol for the machine learning analyses is detailed in Figure 1.

**Figure 1:**
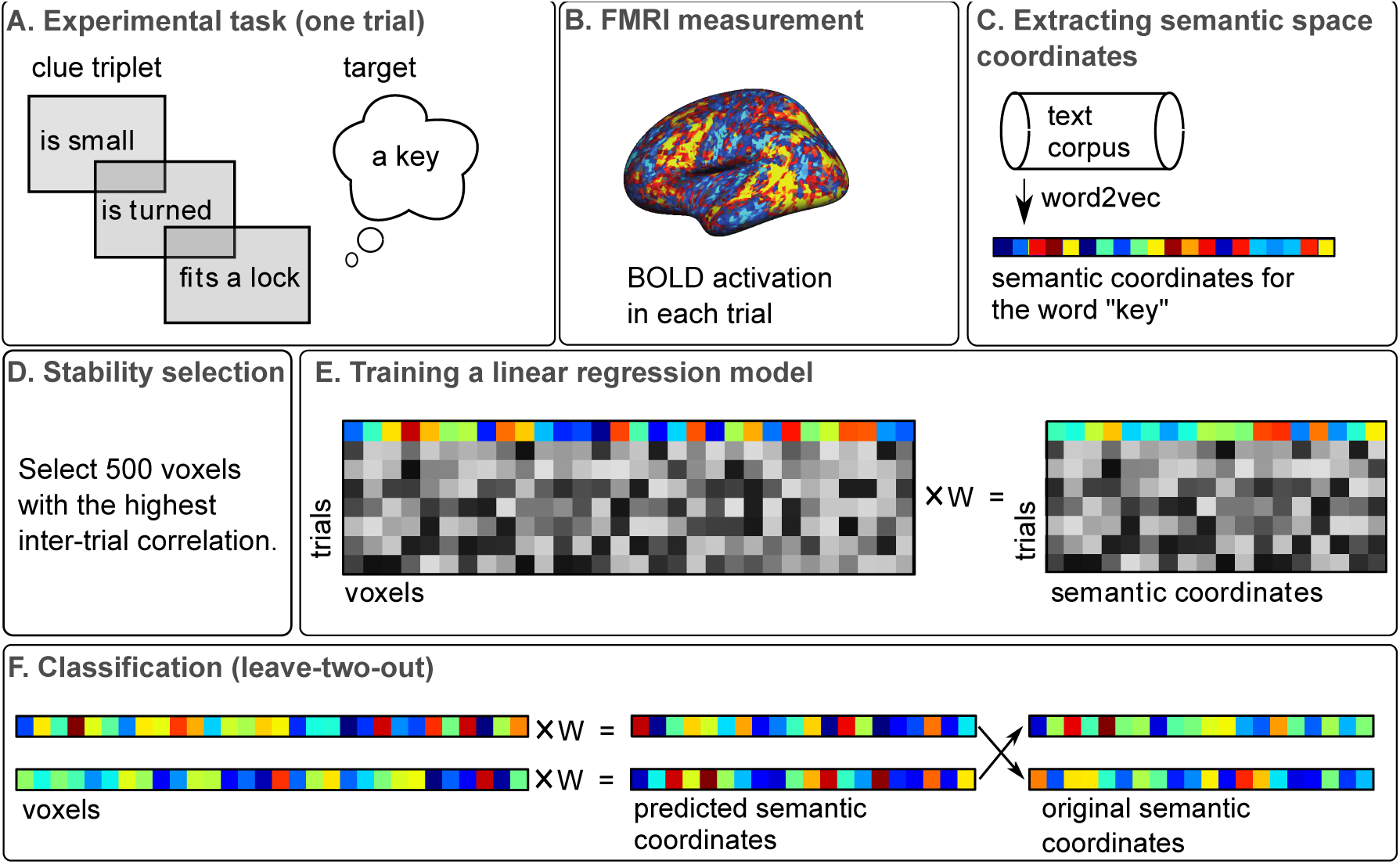
Measurement and analysis protocol. A) Each trial in the experimental task consisted of a clue triplet representing a given target object, e.g., “key”. B) The BOLD activation during each trial was measured. C) Word2vec model was used to extract semantic space coordinates for the implied target word. D) In the stability selection stage, the voxels showing the highest consistency in activation patterns across trials were selected for the analysis. Stability selection was not applied in the single-trial analysis. E) A training set and linear regression were used to map the activation patterns of each target object to its respective semantic coordinates. F) The resulting mapping was evaluated by using it to predict the semantic coordinates of two left-out objects based on BOLD activity. This scheme was repeated for all possible leave-two-out pairs.

The overall level of classification accuracy using the semantic coordinates of the unpresented target objects was high, ranging from 76.1 % to 93.3 % correct classifications across subjects (mean = 87.2 %, SD = 4.9; see also Table S1). The null-distribution for chance level performance was determined through a permutation test in which the relationship of each feature vector and its target was randomized in the training set. This process was iterated 1000 times, each time using a randomly selected participant’s brain activation data. Based on the resulting distribution, decoding accuracies >61.5 % were deemed significantly better than chance (*p* < 0.05); this threshold was exceeded by a comfortable margin for all subjects. Decoding across semantic category (e.g., *elephant* vs. *car*) was expectedly more accurate (mean = 94.2 %, SD = 5.0; *p* < 0.05) than decoding within a semantic category (e.g., *elephant* vs. *giraffe*; mean = 64.5 %, SD = 7.1; *p* < 0.05). Decoding accuracies across semantic category were significant (*p* < 0.05) in all participants, whereas decoding accuracies within a semantic category were significant in 12 out of 17 participants (see also supplementary Figure S1 for a confusion matrix).

The aforementioned analysis yielded a bi-directional mapping between BOLD activation patterns and the corpus-derived semantic space. This mapping can be used to predict BOLD activation patterns to any number of novel objects in the text corpus based on their semantic space coordinates. In the supplementary online material (https://users.aalto.fi/˜vanvlm1/guessfmri), the recorded BOLD activation patterns for each target object are visualized along with their semantic space coordinates. Furthermore, we included some additional targets to demonstrate how the mapping can be used to predict the BOLD activation patterns for novel targets.

### Semantic representation of the target object is best decoded by summing many of its features

We next examined whether we can demonstrate pattern completion in the BOLD activity patterns. We tested whether the neural representations elicited by the clues are better defined through semantic coordinates obtained as a summation of all available features linked to a given target object, as compared to only using the exact clues presented to the participants, or the semantic coordinates of the target object alone. For this, we used a single-trial model where no averaging was performed across the six repetitions of the same target concept and all voxels in the grey matter were used (i.e. no stability selection).

The brain activation patterns for each trial were predicted using semantic coordinates obtained via different models: (1) the last clue of each triplet (“clue 3”), (2) sum of the three clues of the triplet (“clue 1+2+3”), (3) the target object and (4) sum of the full list of semantic features typically associated with the target object (“all available features”) (see Figure 5). Thus, the last model also included many features that were never presented in the experiment. The best approximation for multitude of semantic features associated with each target object was obtained by using a list of behaviorally produced object features from the Centre for Speech Language and the Brain dataset (henceforth, the CSLB features; Devereux et al., 2014). Using the same procedure as for the target words, we established a semantic coordinate for each clue and each of the newly listed features using the word2vec model Figure 2.

**Figure 2:**
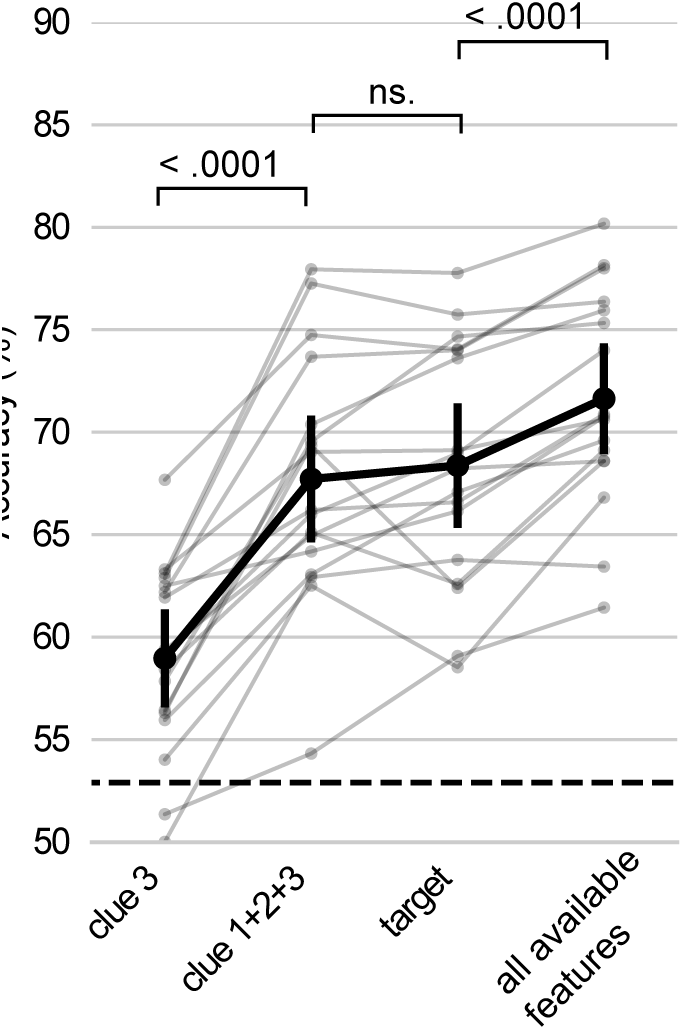
Decoding performance using different models in the singletrial analyses. Each line represents a participant. The mean across participants is indicated as a thick line. Significantly above chance-level decoding performance (*p* < 0.05, based on a permutation test) is marked as a dashed line. The *p*-value of pairwise t-tests (Bonferroni corrected) comparing the accuracy scores between the different models are also indicated.

The best performing model was the one where the resulting semantic coordinates represented the combination of all CSLB features of a target object (Figure 2). This model contains numerous features for a given object: both those included in any one of the trials as well as those not presented to the participant in the whole experiment. This model performed better than the target model and the clue 1+2+3 model, which combined the three clues used to probe the target in a given trial. The next best model was the one using the semantic coordinates of the single target word, on par with the model using the semantic coordinates of the combination of the three presented clues in a given trial. The model using only the last clue of each triplet performed at the lowest level of all.

### Activation patterns in the ventral stream and the perirhinal cortex correlate with the semantic similarity of the full feature sets

In the next step, we aimed to test the neuroanatomical hypothesis of the semantic pattern completion. We used RSA to determine the brain-wide set of regions in which activation patterns were correlated with the semantic similarity structure among the 60 target objects, as represented by the semantic coordinates of all available features (i.e., the best performing model in the zero-shot decoding). The similarity was defined as the cosine distance between the corpus-derived semantic coordinates. For this analysis, we used brain data averaged across the six repetitions of the same target object. Based on the permutation test, the analyses resulted in seven clusters. The largest cluster was centered in the left middle occipital gyrus extending to the middle temporal gyrus, inferior parietal gyrus, superior parietal gyrus, supramarginal gyrus and the rostral extent of the lateral occipital cortex (Table 1, Figure 3). Medially this cluster encompassed parts of the precuneus, isthmus and cingulate gyrus. This large cluster also extended anteriorly to the medial and inferior aspects of the temporal lobes bilaterally and covered a large extent of particularly the left but also the right fusiform gyrus. The cluster extended bilaterally to both PRCs (Table 1, see e.g., Insausti et al., 1998; Kivisaari, Probst, et al., 2013). The locations of all peak voxels are reported in Table 1.

**Table 1:** Peaks of the significant RSA searchlight clusters (*k* > 10)

| Anatomical location | Pseudo-$t$ | Cluster extent | Voxel-level $p$ (FDR) | $x$ | $y$ | $z$ |
| --- | --- | --- | --- | --- | --- | --- |
| Left occipital middle | 10.97 | 10104 | 0.0008 | -41 | -78 | 24 |
| Left inferior frontal | 5.16 | 686 | 0.0008 | 47 | 38 | 15 |
| Vermis | 3.82 | 51 | 0.0052 | 0 | -59 | -35 |
| Left superior frontal | 3.56 | 14 | 0.0048 | -19 | 63 | 18 |
| Right inferior temporal | 3.55 | 17 | 0.0076 | 62 | -15 | -25 |
| Right inferior parietal | 3.41 | 20 | 0.0088 | 37 | -40 | 46 |
| Left frontal superior medial | 3.14 | 13 | 0.0079 | 0 | 53 | 43 |

Effects in clusters smaller than 10 voxels are not shown. FDR = False Discovery Rate.

**Figure 3:**
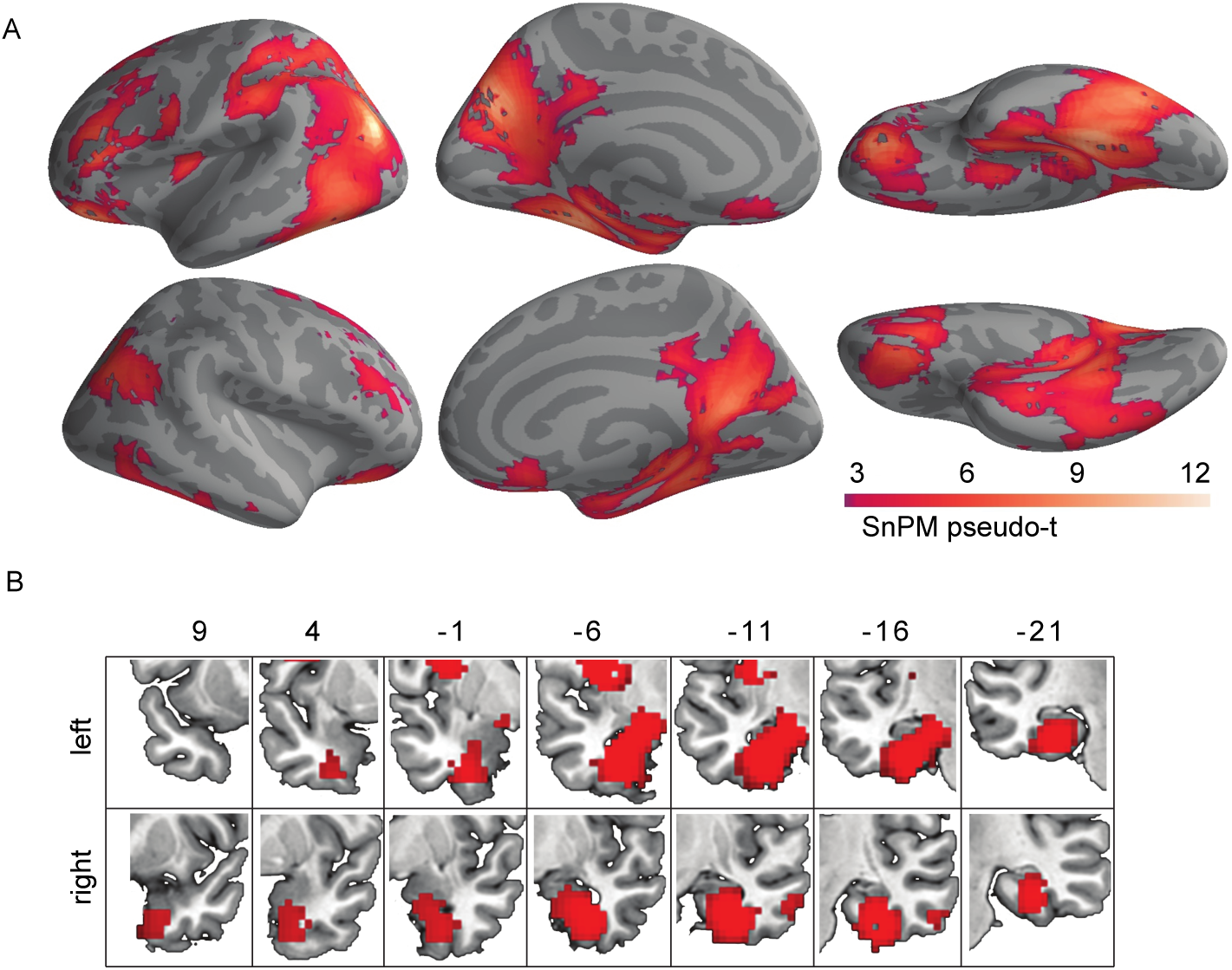
RSA searchlight results. A) Brain regions whose activation patterns correlated with the semantic similarity of the target objects when they were each represented by a combination of their full semantic feature set. B) A coronal view of the anterior extent of the left and right temporal lobes. The clusters overlap with the bilateral PRC.

## Discussion

Humans are able to recognize objects and understand their rich meanings even when only limited information about them is available. In this study, we simulated such a situation by presenting the participants with brief verbal descriptions of sixty objects and asking them to guess the identity of each of them. We showed that it was possible to decode the implied target object with high accuracy without ever showing the object explicitly suggesting that the clues triggered a coherent representation of the target object. The single-trial results further demonstrated that the brain activation patterns elicited by the guessing game paradigm indeed contained more information about each target object than what was initially given as input in the experiment. This suggests that the entire neural representation of an object became activated based on partial stimulation in the form of only few features. This provides neuroimaging evidence on semantic pattern completion whereby limited information in the environment is used to reconstruct coherent object representations.

Distributed accounts of semantic representations postulate that neural representations of objects can be modeled as unique and consistent distributions of activity across a set of perceptual and semantic feature nodes (e.g., Farah and McClelland, 1991; McRae, Sa, et al., 1997; Tyler, Moss, et al., 2000; Tyler and Moss, 2001). This model has been successful in describing, not only the healthy semantic system (e.g., Masson, 1995; McRae, Cree, et al., 2005), but also patterns of semantic impairments associated with brain damage (Kivisaari, Tyler, et al., 2012; Moss et al., 2002; Plaut and Shallice, 1993). Importantly, such a feature-based distributed system also gives an account on how information is reconstructed from incomplete patterns of information. Specifically, the activation in a subset of feature nodes is postulated to propagate in the network based on connection weights which, in turn, are based on experience on co-occurence (Masson, 1995; McClelland and Rogers, 2003; Rumelhart et al., 1986).

We found evidence of semantic pattern completion by combining neuroimaging and machine learning with corpus-derived coordinates of objects and their features in a shared semantic space. Using single trials, we found the best mapping using semantic coordinates created by summing many features for each given object, including features never presented to the participant. This model performed significantly better than the model using the semantic coordinates for the target object alone or that using the sum of the clues presented to the participant in the given trial. This finding therefore provides neuroimaging markers of pattern completion, that is, that activation in a subset of object features leads to activation in a network of features associated with a given object entity (cf. Masson, 1995; McClelland and Rogers, 2003; Plaut and Shallice, 1993; Rumelhart et al., 1986).

A combination of the present RSA results and previous research on visual processing may provide insights on how such semantic pattern completion of objects takes place in the brain. Studies on visual object recognition suggest that pattern completion in the visual domain takes place in the ventral stream via recurrent connections (Tang et al., 2014; Clarke, Taylor, et al., 2011). The importance of the ventral stream in processing also the meaning of visual objects has been demonstrated by Clarke and Tyler, (2014). In that study, the authors presented participants with a large set of naturalistic color photographs and showed that regions in the lateral occipital cortex and ventral stream were sensitive to the semantic similarity of the presented visual objects. The anatomical pattern of the RSA results in the ventral stream in our study bears remarkable similarity to those of Clarke and Tyler, (2014) despite the fact that we never showed images or pictorial stimuli to our participants. It is possible that reading descriptions about objects, such as in the guessing gametask, recruits embodied visual representations of objects and, thus, recruit the ventral stream system (see also Anderson et al., 2015). Our results suggest that this system may also play a role in pattern completion of meaningful object representations.

The significant link between brain activation patterns and the similarity of semantic features of the objects was observed at a very high level of ventral stream hierarchy. In fact, the cluster that showed sensitivity to the semantic similarity of the target objects extended all the way anteriorly to the PRC which is located at the apex of this system. Importantly, this region has been highlighted in item-specific processing of object meaning using a visual object naming task (Clarke and Tyler, 2014) and a property-verification task with pictures and words (Bruffaerts et al., 2013). Our findings extend these results by showing that this region is involved in item-level processing of objects even in the absence of pictorial stimuli. Moreover, the findings corroborate those by Taylor, Moss, et al., (2006) and Taylor, Stamatakis, et al., (2009) who showed that the PRC is involved in binding features from multiple modalities. Importantly, the current study demonstrates that these features need not be visual or auditory but they may also come in the form of more abstract semantic properties. Therefore, this study strongly supports the claim that this region is involved in processing object meaning.

We found a set of other regions that were associated with the semantic similarity of the target objects in addition to those in the ventral stream. These regions include the temporo-parietal junction and inferior frontal cortex, whose involvement may reflect the verbal nature of the task and conceptual and lexical preparation for the verbal response (Indefrey and Levelt, 2004). Other regions include the bilateral retrosplenial cortex, which in previous research has been associated with visual imagery and memory, and whose involvement can partly be explained by specific strategies employed in the task (for a review, see Vann et al., 2009). Importantly, the network of areas revealed by the RSA analysis are also likely to be a reflection of the distributed nature of the semantic representations themselves. Indeed, Huth, Nishimoto, et al., (2012) and Huth, Heer, et al., (2016) showed that semantic information in the brain is organized systematically as smooth gradients reflecting semantic similarity in wide-spread and distributed regions of the brain. Therefore, we postulate that these regions are relevant in presenting concrete objects such as those targeted in the current experiment.

Using stability selection and data averaged across all six trials of the same target object resulted in a high decoding accuracy that was comparable to those in studies which have used colored photographs as stimuli (e.g., Mitchell et al., 2008). In the past, semantic category-level decoding performance has, at least partly, been attributed to a robust response to low-level visual features (Rice et al., 2014). However, the present results demonstrate that these visual attributes are not necessarily needed in order to achieve high-level decoding accuracy. Moreover, we suggest that the guessing game paradigm used in this study is highly engaging from the participant’s point of view, leading to elaborate processing of the target stimuli. Therefore, we suggest that it is particularly well-suited for experimentally accessing semantic representations.

The present neuroimaging study used a novel experimental design to examine how the brain completes patterns of fragmented information into meaningful, coherent semantic representations. This design, coupled with our machine learning models, allowed us to study, for the first time, how the brain takes advantage of very limited information and enriches it with prior knowledge of object meaning. The present results give strong support for the distributed, feature-based models of semantics in the brain and suggest that the ventral stream is involved in binding the features together into coherent object representations.

## Methods

### Subjects

18 native Finnish-speaking, right-handed individuals with no history of developmental or acquired language or other neurological disorders participated in the study. The participants were recruited via student mailing lists in the Aalto University. One participant chose not to finish all measurement runs and was therefore excluded from data analysis. Thus, the sample consisted of 17 individuals (mean age = 21.1 years; SD = 3.4 years; mean education = 12.8 years, SD = 1.7 years; 10 identified themselves as females, and seven as males). All of the participants gave a written informed consent before participating in this study. The study was approved by the Aalto University Research Ethics Committee.

### Stimuli

The stimuli consisted of 540 brief verbal descriptions of 60 target objects in Finnish (9 to 29 characters including spaces, mean 17.5, SD 3.6). Fifty-eight target objects were selected from the CSLB property data set (Devereux et al., 2014). We additionally included two target objects that were not part of the CSLB data [forklift (Finnish: “trukki”) and metro (subway) (Finnish: “metro”)]. One fourth (*n* = 15) of the target objects fell into each of the following semantic categories: animal, fruit/vegetable, tool and vehicle. We created nine clues (i.e., descriptions) per each target object by translating and adapting semantic features from the CSLB data. For the two objects not included in the CSLB data set, we selected six features from that set that applied to the target object and additionally created three new highly distinctive features. We also created 29 new clues (5.3 % of clues in total) in cases where sufficiently many suitable clues were not available in the CSLB data set. The first, second and third clues were matched on length across the four semantic categories (*p* > 0.59 for all).

The nine clues assigned to each target object were further divided into three clue triplets. When feasible, the presentation order of the clues within a triplet was sorted such that the first clue in each triplet was the least distinctive (e.g., ‘has four legs’), and the following two clues increasingly distinctive (e.g., “is found in the savannah” > “has a trunk”) based on the CSLB feature norm data (Devereux et al., 2014). The purpose of this approach was to ensure that the participants would guess the target object approximately at the same stage (i.e., at the third clue).

Each individual clue was repeated twice in the fMRI experiment, once in Set 1 and once in Set 2, with the two sets presented on different days. The clue combinations were rearranged such that each clue’s position in a triplet was retained, i.e., the first clue of the triplet in Set 1 was always the first clue of a triplet in Set 2, but the clues it was grouped with were not identical in both sets. This procedure resulted in six unique clue triplets for each target object which were presented in six separate blocks. The order of sets (across measurement days) and blocks (within a set) was balanced across subjects. The full list of clues (Set 1) can be found in https://github.com/AaltoImagingLanguage/kivisaari_2018.git.

### Procedure

The fMRI experiment was conducted on two days, with three measurement sessions (i.e., blocks) on each day. The two measurement days were on average 10 days apart (mean = 9.9, SD = 7.9). Each trial started with a fixation cross (‘+’, duration: 300 ms) after which the clues were presented one after another. The clue duration was 1000 ms and the first two clues were followed by a blank screen for 200 ms. The third clue was followed by a jittered interval (mean = 8.0 s, min = 4.0 s, max = 11.8 s), after which a string of hash characters “#################” was presented for 1000 ms, prompting the participant to overtly name the target object (Figure 4). The interval between the final clue and the naming prompt was relatively long as we attempted to minimize the overlap between the peaks of the BOLD signals. The naming condition was followed by a jittered interval (mean = 4.0 s, min = 2.3 s, max = 6.2 s) after which the next trial started. The jittering was generated using efMRI version 9 (Chris Rorden, Columbia, SC, USA, www.mricro.com). The black text stimuli were presented on a gray background. There were two 18 s rest periods in each measurement session. The rest trials were signaled by a pair of hyphens “--” that the participant was asked to fixate while remaining still.

**Figure 4:**
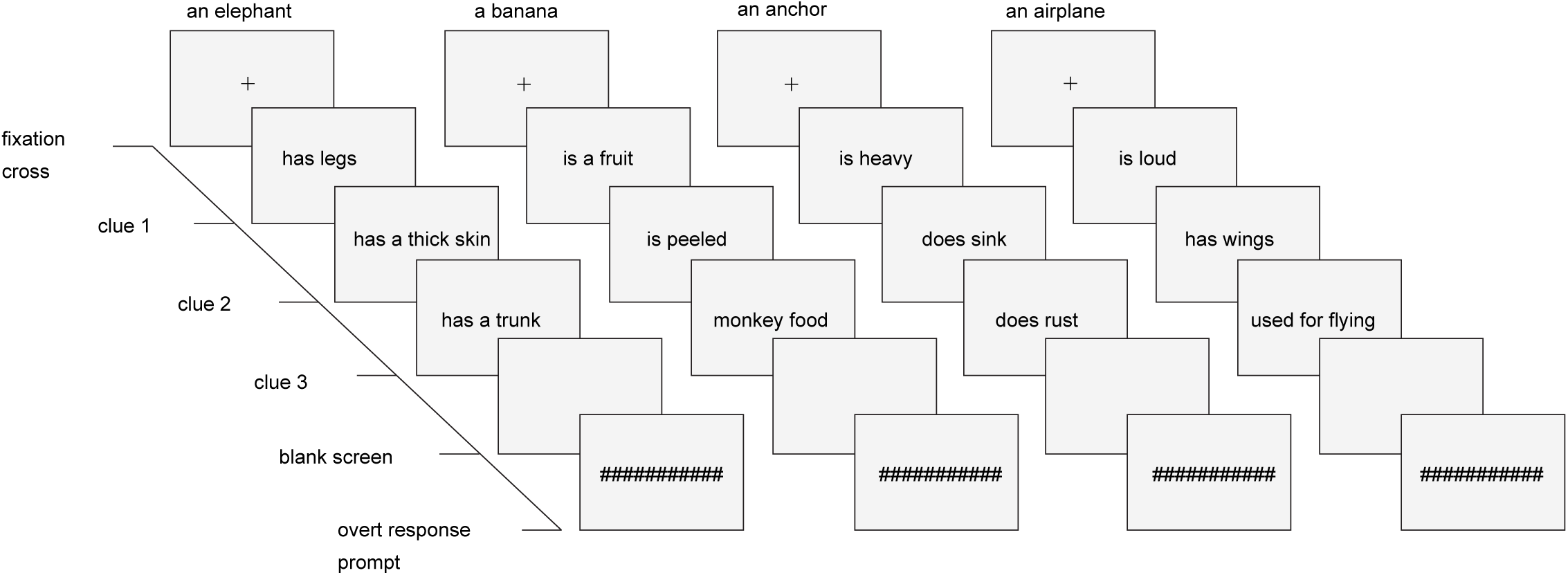
Examples of stimuli and experimental design in fMRI. Three clues were shown one at a time, after which the participants were asked to guess which object they describe (i.e., an elephant, banana, anchor and airplane, respectively). A string of hash characters prompted the participant to utter the name of the target object. The target object itself was never presented to the participants before or during the experiment, either pictorially or as a word, and no feedback regarding correct or incorrect answer was provided.

**Figure 5:**
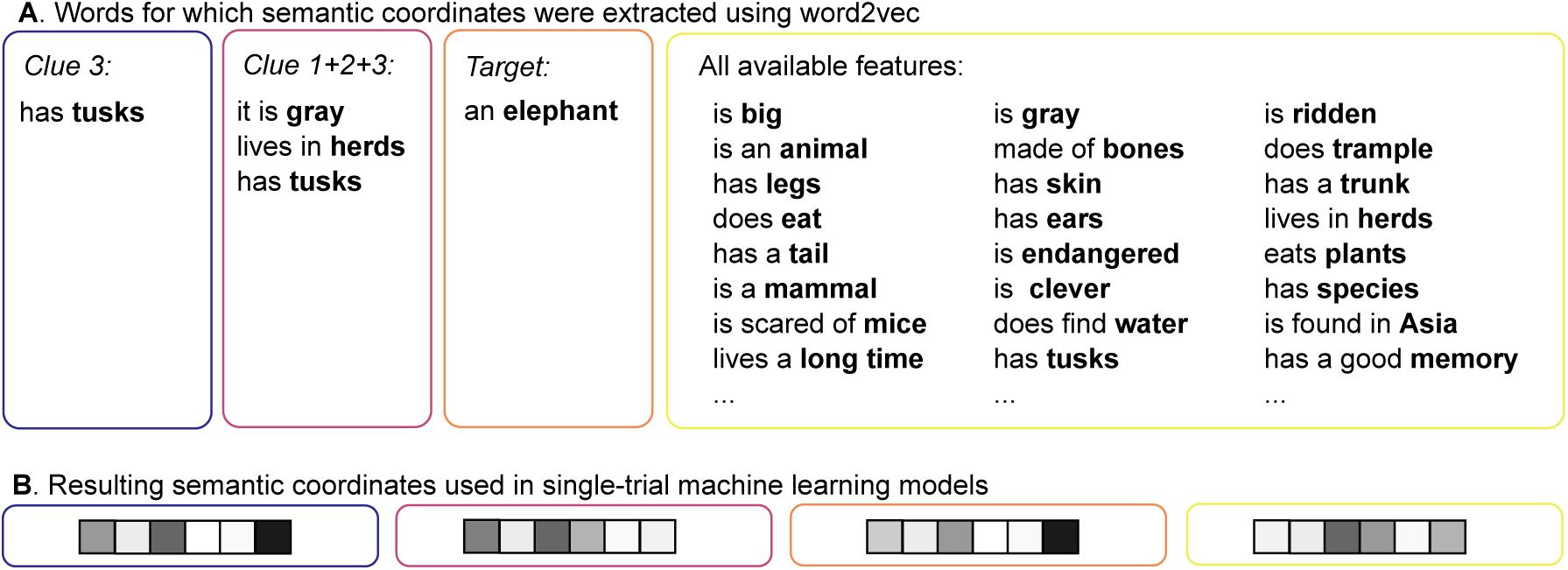
The four types of semantic coordinates used in the single-trial machine learning analysis. The key word whose semantic coordinates was built using word2vec is shown in boldface. The semantic coordinate was either based on one word (i.e., Clue 3 and Target) or several words (i.e., Clue 1+2+3 and All available clues) in which case the final semantic coordinate was a sum of the semantic coordinates of all words in the respective model.

### Functional MRI data acquisition

Participants were scanned with a Siemens 3T Skyra Magnetom MRI device using a custom 30-channel receiver head-coil. We acquired echo-planar imaging (EPI) volumes in axial oblique angle using an acquisition matrix of 64 × 64 with 3.1 mm 3× 1 mm voxel dimensions. The following acquisition parameters were used: TE = 32 ms, TR = 2.4 s, flip angle = 90°, slices = 41, FOV = 200 mm, phase resolution = 100 %. A structural T1-weighted MPRAGE volume was also acquired (TE = 3.3 ms, TR = 1.1 s, slices = 176, FOV = 256 mm, phase resolution = 100 %).

The stimuli were controlled using Presentation^®^ 15.0 software (www.neurobs.com) running on a Dell Optiplex 960 PC. The stimuli were projected to a mirror mounted on the head-coil using a Panasonic PTDZ110XEJ projector using 1920 × 1200 resolution and 60 Hz frequency. Participants’ verbal responses were recorded using an OptoAcoustics (Or-Yehuda, Israel) FOMRI-III optic microphone with OptoActive noise control. The microphone was mounted on the head-coil.

### Semantic space from text corpus data

The model of semantic space used in the decoding was estimated from a 1.5 billion token Internet-derived text corpus in lemmatized Finnish (Kanerva and Ginter, 2014). The semantic space was built using a word2vec skip-gram model with a maximum context of 5 + 5 words (5 words before and after the target word) (Kanerva and Ginter, 2014). The skipgram model is a fast and efficient method for learning dense vector representations of words from large amounts of unstructured text data. The model objective is to find vector representations that are useful for predicting the surrounding words in a sentence given a target word (Mikolov, Chen, et al., 2013; Mikolov, Sutskever, et al., 2013). The code is available online at https://code.google.com/archive/p/word2vec, and the word vector data set used is available online at http://bionlp-www.utu.fi/fin-vector-space-models/fin-word2vec-lemma.bin. The word vectors of the model have the dimensionality of 300, and they were used in the machine learning analyses and the representational similarity analysis (RSA; Kriegeskorte et al., 2008). Note that single dimensions of the semantic space are not interpretable.

We extracted the semantic coordinates for each target object word, which resulted in a 60 × 300 matrix (i.e., number of target stimuli × number of dimensions of the semantic space). Furthermore, we extracted semantic coordinates from the word2vec model for all clues and all the individual features of each target item as provided in the behavioral CSLB norm data. We used altogether four sets of semantic space coordinates: (1) the last single clue of the triplet that was used as the onset for the fMRI response (Clue 3); (2) the sum of the first, second and third clue of the triplet that were used to probe a given target word (Clue 1+2+3), (3) the target word alone (Target) and (4) the sum of the semantic coordinates of all features for a given target object available in the CSLB data set (All available features; Devereux et al., 2014), including features that were never presented to the participant. In cases where the clue/feature consisted of more than one word, we selected and lemmatized one key word (e.g., has legs → leg) and extracted the corresponding semantic coordinate from the corpus data.

### FMRI data preprocessing

The preprocessing was performed using SPM8 software (Wellcome Trust Centre for Neuroimaging, University College London, UK) running on Matlab (MATLAB 2014a, MathWorks In., Natic, MA). The EPI volumes were first corrected for slice timing and head motion and coregistered to the structural volume of the same participant. We used a General Linear Model approach, where the model contained the head motion and session parameters as nuisance regressors as well as high-pass filtering. Each of the target objects in each of the six blocks was modeled by convolving a canonical hemodynamic response function from the onset of the last clue of a triplet. All analyses were run on native-space unsmoothed data. For visualization purposes, the data was co-registered to Montreal Neurological Institute (MNI) reference space (Talairach & Tournoux, 1988). Anatomical labeling was based on the AAL atlas (Tzourio-Mazoyer et al., 2002) unless otherwise cited.

### Zero-shot learning and model evaluation

The machine learning analyses were run on Python 3 (www.python.org) using the Anaconda distribution (2016) and the scikit-learn module (Pedregosa et al., 2011). The machine learning model implemented in this study evaluated the contributions of the brain activation patterns to each of the 300 features of the corpus (Figure 1). The model was trained by using a subset (*n* - 2) of the items and the respective multi-dimensional semantic coordinates such that, in the end, each semantic dimension was associated with a particular weighted activation pattern. For this, we used multiple regression with regularization parameters. The model was evaluated after the training such that the predicted semantic coordinates of the two left-out objects were compared with the original corpus-derived (“true”) semantic coordinates. The classification outcome was determined using cosine distance. This training and evaluation process was iterated 1770 times to cover all leave-two-out combinations. We evaluated the level of statistical significance using a permutation test with 1000 iterations, randomly selected subjects and randomly shuffled order of the semantic coordinates across the target objects.

### Analyses on averaged data and stability selection

We focused the machine learning analyses on a subset of voxels that showed a consistent activation pattern across the six trials of each target object (Just et al., 2010; Mitchell et al., 2008). First, we masked the native space beta images using an individual gray matter mask extracted from the SPM segmentation. We then extracted beta values for each voxel of each repeated trial (*n* = 6) of each object (*n* = 58, i.e., excluding the leave-two-out objects at each iteration). We then calculated pairwise Pearson correlations across the six repetitions of each target object and averaged the correlations over the 58 target objects in the training set. Finally, the 500 most stable voxels, i.e., those with the highest average correlation, were selected for further analyses.

### Single-trial analysis

In the single-trial analysis, no averaging was performed over the six trials of the same target object, but each trial was considered as an isolated event. The brain activation patterns related to each trial were then used to predict the target object or different sets of features (for details, please see section: Semantic space from text corpus data). In the classification stage, we ignored those pairwise leave-two-out combinations that would have included the same target object from different trials (e.g., dog block 1 vs. dog block 2, hammer block 2 vs. hammer block 4). The performance of different models was compared using a pairwise *t*-test using a Bonferroni correction. Note that we did not use stability selection in the single-trial analysis, since there were no repeated trials over which stability selection could sensibly have been performed. Furthermore, as each trial had a different set of clues, we did not want to potentially wipe out this variability.

### Visualization of the zero-shot results

To demonstrate the mapping between the brain and semantic space learned by the zero-shot decoding algorithm, we have created an interactive visualization (https://users.aalto.fi/∼vanvlm1/guessfmri) that shows for each target object, its coordinates in the semantic space and the corresponding BOLD activation pattern, averaged across the six trials. T-Distributed Stochastic Neighbor Embedding (t-SNE) (van der Maaten and Hinton, 2008) was used to obtain a two-dimensional visualization of the semantic space and pycortex (Gao et al. 2015) was used to visualize the BOLD activation pattern. To illustrate that the mapping between the brain and semantic space is defined at all coordinates, we added 19 new targets (*mouse, parrot, chicken, goat, lynx, peach, grapefruit, beetroot, broccoli, lettuce, plane, screw, plate, watch, tape, tram, tank, dinghy, gondola*) to the interactive visualization. By reversing the mapping to obtain a linear transformation between the semantic space and the brain (Haufe et al., 2014), BOLD activation patterns were predicted for these novel items.

### Representational similarity analysis

In the RSA analysis, we used data averaged over the six repetitions of each target object so as to maximize the signal-to-noise-ratio. We used searchlight mapping (Kriegeskorte et al., 2008) and RSA toolbox (Nili et al., 2014) running on Matlab 2014a to find regions where similarity of activation patterns was related to the similarity structure of the all available features of the target objects. A searchlight (radius = 7 mm) was formed around each voxel in the measured volume and the distance (1 – Pearson’s correlation) was computed between each averaged trial. This procedure yielded a symmetrical 60 × 60 BOLD representational dissimilarity matrix (RDM), where the value in each cell reflects the dissimilarity of activation patterns between a pair of target objects. The resulting RDM was compared to that derived from the summed semantic coordinates of all available features. This semantic model RDM was also a 60 × 60 matrix, where the value in each voxel represents the cosine distance between these semantic coordinates. The Spearman’s rank correlation of the BOLD RDM and semantic model RDM were Fisher transformed in order to make them normally distributed. Finally, the resulting correlation values were projected back onto each searchlight’s center voxel and subjected to group-level analyses.

The correlation maps of each participant were transformed into MNI space and smoothed at 6 FWHM. The resulting normalized and smoothed images of each participant were subjected to a group-level SnPM analysis using variance smoothing of 6 FWHM and 10000 permutations (SnPM13; http://go.warwick.ac.uk/tenichols/snpm). Clusters surviving familywise-error-corrected *p* < 0.05 that are over 10 voxels in size are reported.

## Data availability statement

Information related to data and software availability is detailed in Supplementary Table S2. The relevant data used in this study are available upon request to the editors and reviewers and researchers who meet the criteria for access to confidential data. Qualified researchers may contact the secretary of the Aalto University Research Ethics Committee, Jari Söderström. Ethical restrictions prevent the authors from making the raw MRI data publicly available, as this would compromise participant privacy and consent. These restrictions have been imposed by the Aalto University Research Ethics Committee, in compliance with Finnish legislation on Data Protection.

## Acknowledgements

We acknowledge the computational resources provided by the Aalto Science-IT project and thank Marita Kattelus for measurement assistance. We also thank Filip Ginter and Jenna Kanerva from the Turku BioNLP Group, University of Turku for providing the Finnish corpus-based model. We also thank Hanna Renvall, Jan Kujala, Mia Liljeströn and Anni Nora for insights and helpful discussions. This research was funded by the Academy of Finland (grant #286070 to S.L.K., #310988 to M.v.V, #287474 to A.H., #255349, #256459 and #283071 to R.S.), the Aalto Brain Center (M.v.V and T.L-K.), and the Sigrid Jusélius Foundation (to R.S.).

## Author contributions

Conceptualization, S.L.K, A.H., R.S.; Methodology, S.L.K, M.v.V., R.S., A.F., T.L.K.; Investigation, S.L.K.; Interpretation: S.L.K., M.v.V., A.H., R.S., Writing – Original Draft, S.L.K.; Writing – Review & Editing, S.L.K, M.v.V., A.F., A.H., T.L.K., R.S.; Funding Acquisition, S.L.K, R.S.

## Competing interests

The authors declare no competing interests.

